# Cellular resolution imaging of vestibular processing across the larval zebrafish brain

**DOI:** 10.1101/302752

**Authors:** Itia A. Favre-Bulle, Gilles Vanwalleghem, Michael A. Taylor, Halina Rubinsztein-Dunlop, Ethan K. Scott

## Abstract

The vestibular system, which reports on motion and gravity, is essential to postural control, balance, and egocentric representations of movement and space. The motion needed to stimulate the vestibular system complicates studying its circuitry, so we previously developed a method for fictive vestibular stimulation in zebrafish, using optical trapping to apply physical forces to the otoliths. Here, we combine this fictive stimulation with whole-brain calcium imaging at cellular resolution, delivering a comprehensive map of the brain regions and cellular responses involved in basic vestibular processing. We find these responses to be broadly distributed across the brain, with unique profiles of cellular responses and topography in each brain region. The most widespread and abundant responses involve excitation that is rate coded to the stimulus strength. Other responses, localized to the telencephalon and habenulae, show excitation that is only weakly rate coded and that is sensitive to weak stimuli. Finally, numerous brain regions contain neurons that are inhibited by vestibular stimuli, and these inhibited neurons are often tightly localised spatially within their regions. By exerting separate control over the left and right otoliths, we explore the laterality of brain-wide vestibular processing, distinguishing between neurons with unilateral and bilateral vestibular sensitivity, and revealing patterns by which conflicting vestibular signals from the two ears can be mutually cancelling. Our results show a broader and more extensive network of vestibular responsive neurons than has previously been described in larval zebrafish, and provides a framework for more targeted studies of the underlying functional circuits.

## Introduction

The vestibular system of the inner ear reports on linear acceleration, rotation, and gravity, and plays a critical role in animals’ sense of position and movement through space[1]. Vestibular perception relies on the semicircular canals, which detect rotational acceleration, and otoliths, which report on linear acceleration. These distinct types of information are integrated to produce a coherent sense of the head’s position, movements, and relationship with gravity. The sensory circuitry underlying this perception and integration has been the topic of intense research, largely in primates. Electrophysiological studies have shown responses to vestibular stimuli across several brain regions, including in the vestibular nuclei[2], the thalamus[3, 4], various cortical regions[5, 6], and the cerebellum[7, 8]. A host of electrophysiological, lesion, tract tracing, and electroencephalography studies support the idea of a vestibulo-thalamo-cortical stream of vestibular information with extensive crosstalk to the cerebellum (reviewed in [9]). Within this framework, however, many of the cellular microcircuits mediating vestibular processing remain uncharacterised. This is due in part to the complexity of this system, which needs to reconcile and integrate separate streams of information from the otoliths, the semicircular canals, and other modalities[10], but it also springs from a fundamental obstacle to studying the vestibular system: vestibular stimulation requires that the subject be rotated or accelerated. This fact complicates popular approaches for studying neural function, such as electrophysiology and functional imaging.

Recently, we developed a technique based in optical trapping (OT) for the fictive stimulation of the vestibular system in larval zebrafish[11]. This study showed that a focused infrared laser, targeted at the edge of the utricular otolith, could apply a physical force to the otolith. The resulting movement of the otolith was sufficient to stimulate the vestibular system, driving tail and eye movements predicted to compensate for the perceived acceleration and roll. The goals for this approach were to develop a stationary platform in which to study vestibular processing, and to achieve the new ability to stimulate subsets of the vestibular structures (left and right, for instance) in isolation.

Given the external development and optical transparency of zebrafish larvae, this opens the door to calcium imaging of neurons involved in vestibular processing[12]. The introduction of genetically encoded calcium indicators (GECIs) has allowed the observation of activity across large populations of neurons with single-cell resolution, using fluorescence techniques such as 2-photon or selective plane illumination microscopy (SPIM). For sensory modalities that are readily compatible with stationary preparations, such as olfaction[13], audition[14-16], and especially vision[17-20], such studies have uncovered patterns of activity spanning thousands of neurons, in some cases encompassing the entire brain.

In the work reported here, we have incorporated optical trapping into a customized SPIM microscope, allowing us to deliver fictive vestibular stimuli to zebrafish larvae while performing calcium imaging. We report individual neurons’ responses to a range of stimulus strengths, identifying two functional classes of neurons that are excited by vestibular stimuli and one class that is inhibited. We identify ten brain regions with consistent vestibular responses, and for each, provide a functional profile of the registered anatomical locations, response types, and laterality of their constituent vestibular neurons. We also report on the distinct cellular responses that are elicited by fictive vestibular stimuli oriented in different directions. This provides, to our knowledge, the first calcium imaging performed on vestibular processing, and the first brain-wide map of vestibular processing at cellular resolution.

## Results

### A custom setup for whole-brain calcium imaging during fictive vestibular stimulation

We have previously shown that optical traps, applied to the utricular otoliths of larval zebrafish, can produce forces that simulate motion and elicit appropriate behavioral responses[11]. First, we sought to reproduce these results in an optical setup compatible with SPIM imaging of the GECI GCaMP6s. Our microscope setup (Figure 1A) combines OT, two scanning light sheets, a fluorescence emission channel, and a camera for behavioral imaging. For OT, we produced two 1064nm beams (one for each otolith), each of which can be targeted individually using gimbal-mounted mirrors. We generated scanning light sheets using galvanometric (galvo) mirrors, and detected fluorescent emissions through the same objective that delivered the OT. We adjusted the imaging focal plane with an electrically tuneable lens (ETL), synchronized with the galvo mirrors, thus permitting us to perform volumetric imaging without moving the specimen or the imaging objective (Figure 1B). Targeting of the OT requires sub-micron precision, and as such, this stability was necessary for consistent application of trapping forces. Finally, we imaged movements of the tail through a low-power objective below the larva (camera 1, Figure 1A) and movements of the eyes using the fluorescence imaging camera (camera 2, Figure 1A).

**Figure 1:**
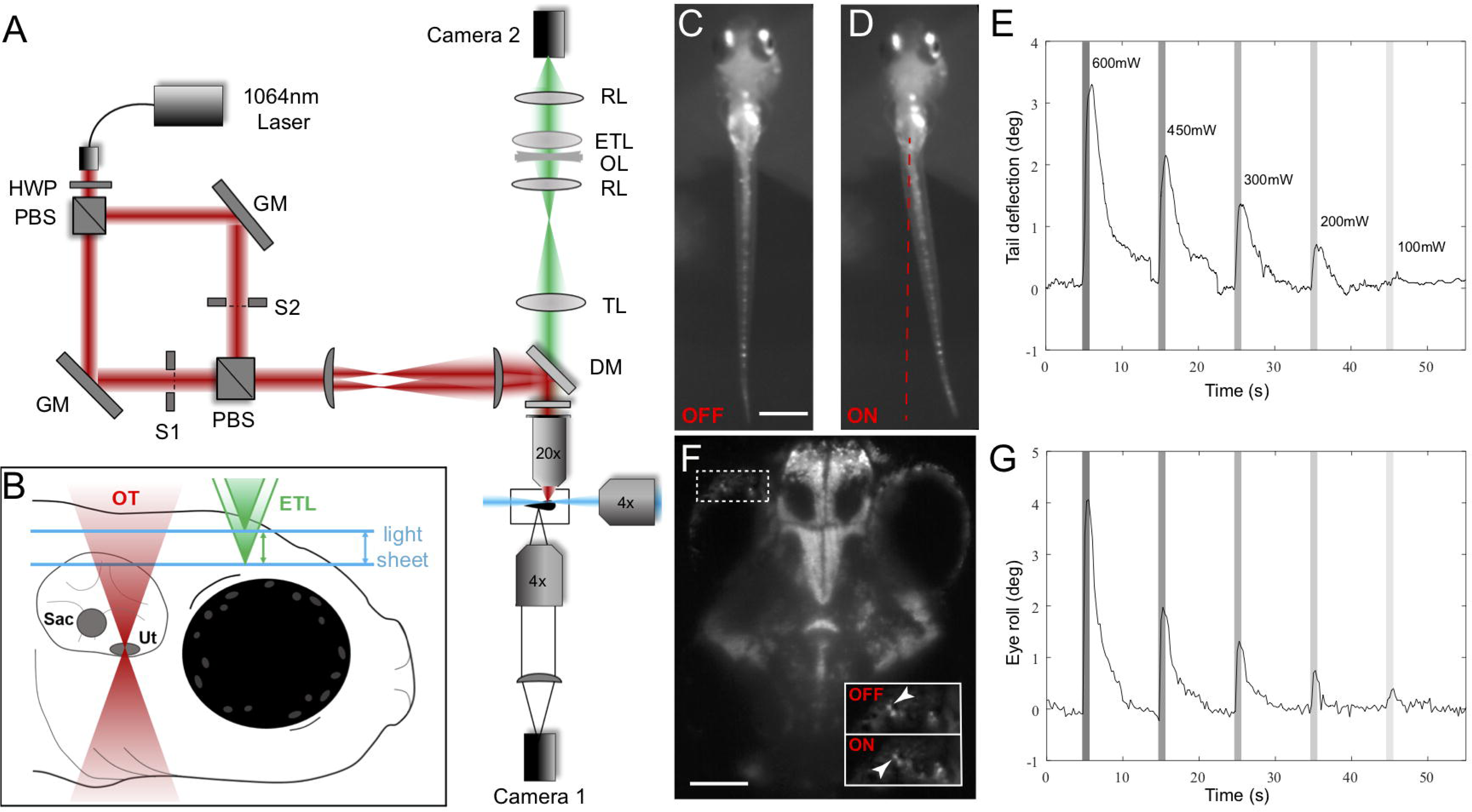
Combining SPIM imaging and OT in larval zebrafish. (A) A conceptual diagram of our OT SPIM microscope. A 1064nm laser beam passed through a half wave plate (HWP) and a polarising beam splitter (PBS) that produced two beams of same intensity. The two beams were separately steered with gimbal-mounted mirrors (GM) before being recombined and projected onto the back focal plane of a 20x objective via two lenses and a shortpass dichroic mirror (DM). Camera 1 recorded tail movements via a 4x objective located below the larva. (B) a diagram of the approach for volumetric SPIM imaging during OT. Two 488nm light sheets (one from the front and one from the side of the larva), were produced and scanned in the z-axis using 2D galvo mirrors. Camera 2 imaged fluorescent emissions from GCaMP6s through a tube lens (TL), relay lenses (RL), an electrically tuneable lens (ETL), and an offset lens (OL). Coordinated control of the galvo mirrors and ETL allowed volumetric scanning of a 300μm deep volume of the brain. (C and D) Images from Camera 1 of a larva without (C) and with (D) a 600mW trap on the lateral side of the left otolith. Average tail deflections resulting from different OT powers are shown in (E). Powers are indicated and maintained through panel G and other subsequent figures. (F) An image from Camera 2 showing the eye position before (yellow dot) and during (cyan dot) the same OT, and average eye movements are shown in (G). Scale bars represent 500μm in (C) and 100μm in (F). n=6 for (E) and (G). A detailed description of the optical setup and its parts can be found in the STAR Methods section.

Consistent with our previous work[11], we found that optical trapping in this setup drove compensatory movements of both the tail and eyes (Fig 1C-G). An outward force applied to either otolith caused a contralateral tail bend that was proportional to the strength of the OT (Figure 1D, E). These forces also produced power-dependent rolling movements in both eyes (Figure 1F, G), representing the vestibulo-ocular reflex (VOR) that compensates for the larva’s perceived rolling motion. These behavioral responses, which are similar to those that we have previously reported in an optically simpler configuration, suggest that we are successfully triggering the vestibular system in our customised SPIM microscope.

### Fictive vestibular stimulation elicits activity broadly across the brain

We next performed brain-wide GCaMP imaging (in the *elav3:nuc*-*GCaMP6s* transgenic line [21], which expresses GCaMP6s in the nuclei of all neurons) during OT to identify brain regions involved in vestibular perception and processing. We imaged 25 planes at 10μm intervals, sampling from a volume 650μm in X, 800μm Y, and 250μm in Z. We used the CaImAn package for morphological segmentation, thus identifying and extracting the fluorescent traces from regions of interest (ROIs) corresponding to individual neurons, as previously described[22, 23], carried out a constrained nonnegative matrix factorization (CNMF) step to demix overlapping fluorescent signals, and deconvolved traces to denoise them. We then used k-means clustering to identify classes of response that different neurons had to the perceived vestibular stimulation. Finally, we spatially registered the brains of all of our larvae against one another, and registered this combined map to the Z-Brain atlas of the 6 day postfertilization (dpf) larval zebrafish brain[24-26]. This approach delivered the brain positions and response properties of 220,000 individual ROIs across 13 larvae, each spatially registered within the brain regions annotated in Z-Brain. Details of the segmentation, clustering, and registration approaches can be found in the STAR Methods.

Our stimulus train comprised optical traps of five intensities: 600, 450, 200, 100, and 50mW, each delivered three times for a duration of one second. Signals from individual ROIs across 13 6dpf larvae showed 4,000 ROIs with clear and consistent responses to the OT. These fell into three broad functional categories: those with strongly strength-dependent responses (green in Figure 2A and throughout this paper), those with relatively flat responses across all powers (cyan), and those that were inhibited as a result of vestibular stimulation (purple). Because we are using calcium imaging as a proxy for the neurons’ electrical activity, it is difficult to infer spiking patterns for neurons in these ROIs, but the positive relationship between trap strength and calcium response for the green and purple clusters suggest that stimulus strength in rate coded in these ROIs.

**Figure 2:**
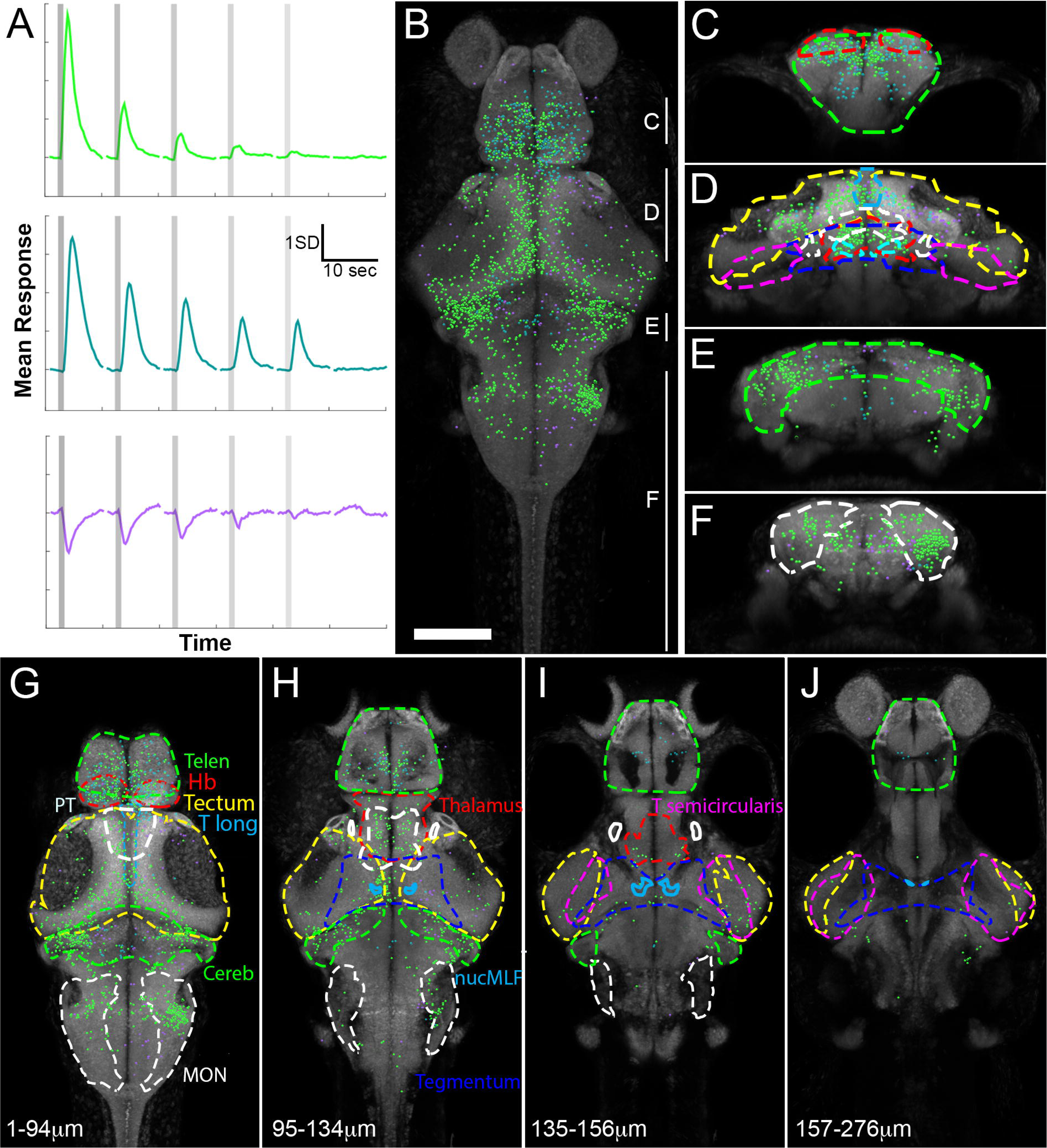
Vestibular responses across the larval zebrafish brain. Functional clustering of 4,000 vestibular-responsive neurons revealed three distinct response profiles (A). This panel shows the mean response across each cluster of neurons to fictive vestibular stimuli at various OT powers (indicated in Figure 1), applied to the lateral edge of the right utricular otolith. Vertical lines show the times of stimulation, and three trials at each power are averaged to produce this graph. (B) A maximum intensity projection of all ROIs’ locations in a combined reference brain (a rotation of this stack is shown in Supplemental Movie 3). ROIs’ colors correspond to their cluster, as shown in (A), and are maintained throughout this paper. (C-F) show coronal virtual stacks at different rostro-caudal positions of the map (rostro-caudal ranges are indicated in B), with responsive brain regions delineated. (G-J) show virtual horizontal stacks at different dorso-ventral positions (indicated distances below the dorsal surface of the brain) with the same regions delineated. Regional outlines are color coded; Telen: telencephalon, Hb: habenulae, PT: pretectum, T Long: torus longitudinalis, Cereb: cerebellum, MON: medial octavolateralis nucleus, nucMLF: nucleus of the medial longitudinal fascicle, T semicircularis: torus semicircularis. Scale bar in B indicates 100μm.

These responsive ROIs were distributed broadly across the brain, with different brain regions containing different mixes of ROIs belonging to each of the three functional clusters (Figure 2 and Supplemental Movie 3). In the forebrain, the telencephalon, the habenulae, the thalamus, and the pretectum each showed consistently responsive neurons. In the midbrain, such responses were found in the tectum, the nucleus of the medial longitudinal fascicle (nMLF), and the tegmentum. Responsive neurons were found throughout the hindbrain, with clear foci in the cerebellum, the medial octavolateral nucleus (MON), and rhombomeres 5-7. Each of these brain regions showed unique properties in terms of the number of responsive ROIs, the functional clusters represented, and the topographical distribution of these responses. In the next section, we will report the responses observed across each of these regions, highlighting the characteristics with implications for functional connectivity and the flow of vestibular information from perception to behavior.

### Vestibular responses are widespread but regionally distinct

In order to ensure we did not miss any minor or underrepresented response types, especially for inhibited neurons that may be missed using CNMF, we performed clustering on the vestibular-responsive ROIs within each region separately, including ROIs that CNMF had discarded. Although there are subtle region-to-region differences in the average responses of these clusters, all clusters essentially matched one of the three described above. This indicates that these are the three cardinal response types to vestibular stimuli in the larval zebrafish brain.

Telencephalic responses to fictive vestibular stimulation (Figure 3A and B) came from two functional clusters: one with significant responses across the spectrum of trap powers, and one that is inhibited as a result of vestibular input (Figure 3C). The former was broadly and evenly distributed across the telencephalon, while inhibited responses were patchy, showing a predominantly ipsilateral distribution with regard to the stimulated ear in the medial pallium (homologous to the amygdala in mammals[27]), and a predominantly contralateral distribution in the lateral pallium (homologous to the hippocampus[27], Figure 3D). The habenulae were notable as the only brain region in which all three clusters were robustly represented (Figure 3D-F). The ROIs for each of these clusters are abundant throughout both habenulae, and there was no obvious spatial pattern, with ROIs from each of the clusters distributed essentially evenly (Figure 3D-E). In the thalamus, a majority of ROIs were excited in a strongly force-dependent manner, but a small proportion of ROIs were inhibited (Figure 3G-I). These inhibited neurons were notably distributed on the ipsilateral side of the thalamus to the stimulus (Figure 3G-H), and were located either in the dorsal thalamus or in the dorsalmost portion of the ventral thalamus (Figure 3H). The fact that the thalamus is not yet nucleated in 6dpf larvae[28] suggests that the distinction between dorsal and ventral thalamus is not clear cut, and may not be of great functional importance. The results nonetheless show that inhibition in the thalamus is predominantly ipsilateral and dorsal within the thalamus as a whole. Similarly in the pretectum (Figure 3J-L), a majority of responsive neurons were excited in a power-dependent manner, although inhibited neurons, mostly in the rostral portion of the pretectum, were also present. The most striking spatial pattern involves ROIs in the migrated pretectal region M1[29, 30] (arrows, Figure 2J and K), where a densely packed group of inhibited ROIs is ipsilateral to the stimulated otolith, with excited responses contralaterally.

**Figure 3:**
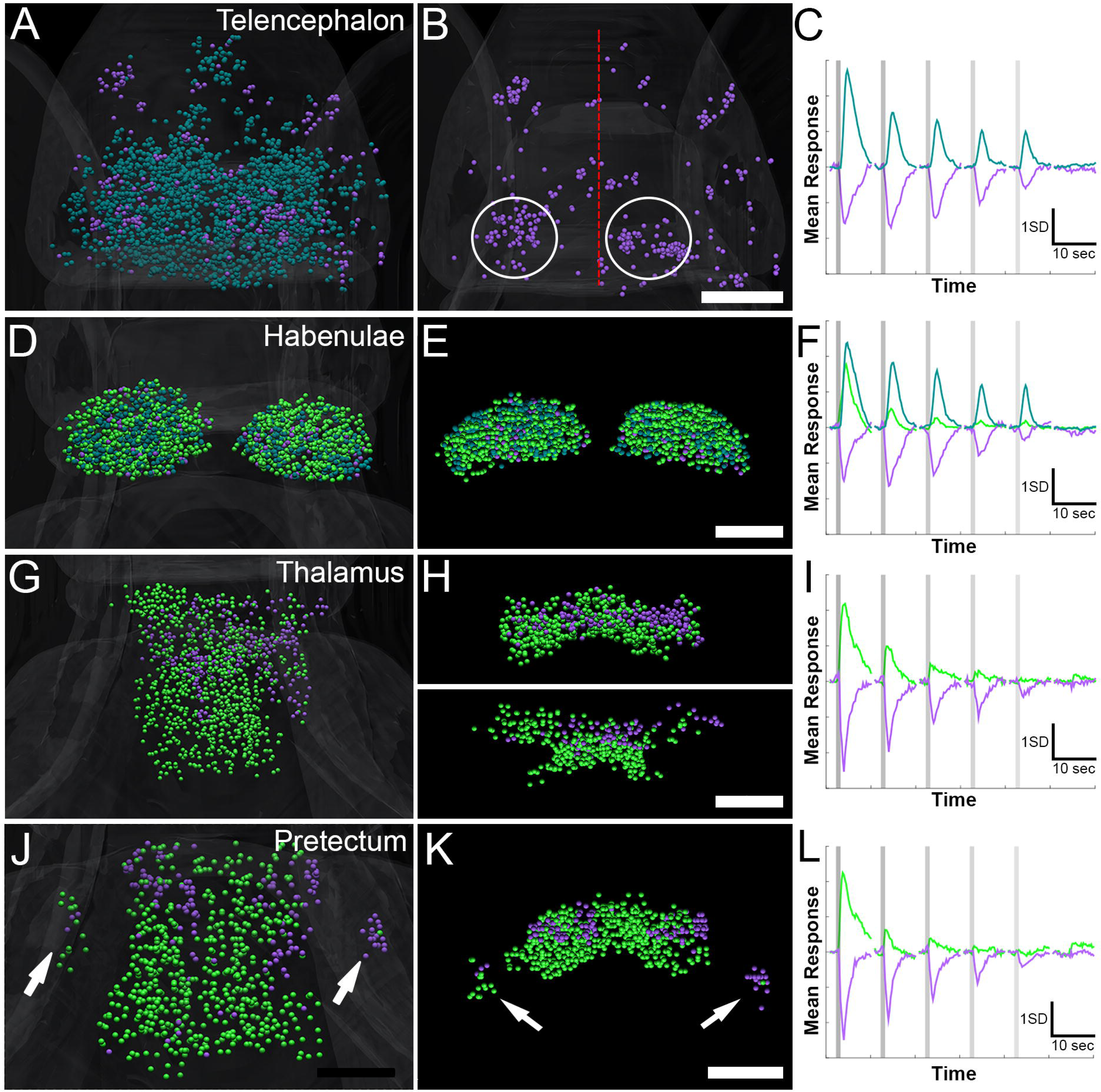
Vestibular responses in the forebrain. Telencephalic responses (dorsal view, A) include excited and inhibited (shown alone in B) ROIs (average traces for these two clusters are shown in C). Several regional concentrations of inhibited ROIs are spread throughout the telencephalon, with a tendency toward medial regions of the ipsilateral (right) and lateral regions of the contralateral (left) side (circles in B, with the midline indicated by a red line). Dorsal (D) and coronal (E) views of the habenulae show even distribution of three clusters (F). Two clusters in the thalamus are shown in dorsal (G) and coronal (H) views, with the dorsal (top, H) and ventral (bottom, H) segmented separately. These clusters’ traces are shown in (I). The pretectum (dorsal in J, and coronal in K) contains ROIs with two similar clusters (L). The laterality of responses is notable in the migrated pretectal region Ml (arrows, K). Scale bars indicate 50μm.

In the midbrain, the tectum is the strongest vestibular-responsive region, showing both strength-tuned excited ROIs and inhibited ROIs (Figure 4A-D), both of which are evident in the periventricular layer (PVL, Figure 4A) and in the neuropil (Figure 4B). While the excited ROIs are roughly evenly distributed, the inhibited ROIs are almost exclusively on the side ipsilateral to the stimulus, and are concentrated in the neuropil, the PVL adjacent to the neuropil, and a concentrated region of the ventral PVL near the midline. The tegmentum had a particularly simple functional profile, with a single cluster of strength-tuned excited ROIs, a majority of which were contralateral to the stimulus (Figure 4E and F). Similarly excited ROIs represented a majority of the relatively few responses found in the nMLF (Figure 4G and H), and again, these ROIs were almost exclusively contralateral to the OT. The small number of inhibited ROIs in the nMLF were exclusively ipsilateral to the OT.

**Figure 4:**
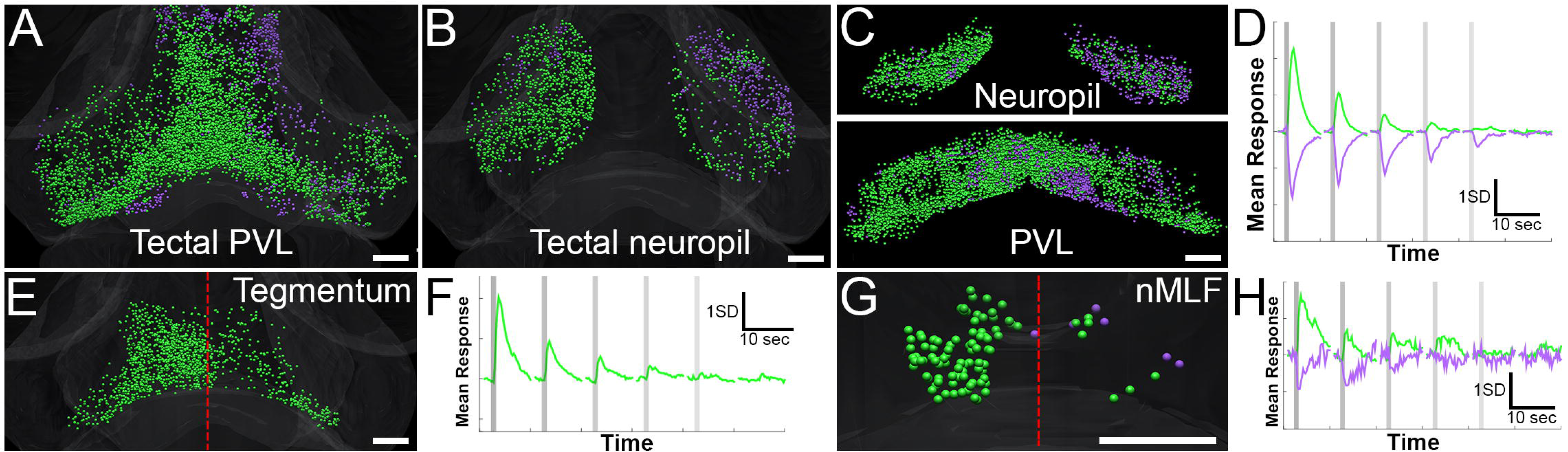
Midbrain responses to vestibular stimuli. Dorsal views of the tectal PVL (A) and neuropil (B), with coronal views (neuropil, top; PVL, bottom) shown in (C). The average responses of these clusters are shown in (D). The tegmentum (dorsal view in E and average response in F) has a single cluster, predominantly contralateral to the right (stimulated) ear. (G and H) The nMLF contains two clusters, with a majority of responsive neurons contralateral to the stimulation. Sparse inhibited ROIs are located ipsilaterally. Red lines in E and G indicate the midline, and scale bars indicate 50μm.

In the hindbrain, the cerebellum is a site of remarkable topography. ROIs responding to strong OT forces are spread across the cerebellum, except for a sharply delineated region in the dorso-medial cerebellum, where almost exclusively inhibited ROIs were observed (Figure 5A-C). Based on their position within the cerebellum, these ROIs are likely to represent Purkinje cells [31, 32]. In the caudal hindbrain, spanning rhombomeres 5-7 (Figure 5D-G), there was a topographical bias, with strength-dependent excited ROIs located roughly equally on the ipsilateral and contralateral sides, and most inhibited ROIs occupying the ipsilateral side in a pattern resembling that already described in the nMLF. In the MON, most responsive ROIs were excited in a strength-dependent manner, and these were more numerous in the ipsilateral MON, adjacent to the stimulated ear (Figure 5H-J). A small number of inhibited ROIs were present exclusively around the periphery of this region, although these may represent neurons from adjacent parts of the hindbrain that were erroneously mapped to the MON during the registration process. This profile for the MON, containing predominantly or exclusively ROIs with power-sensitive excited responses, is consistent with its established role as the conduit between the vestibular ganglion in the ear and the rest of the brain[33].

**Figure 5:**
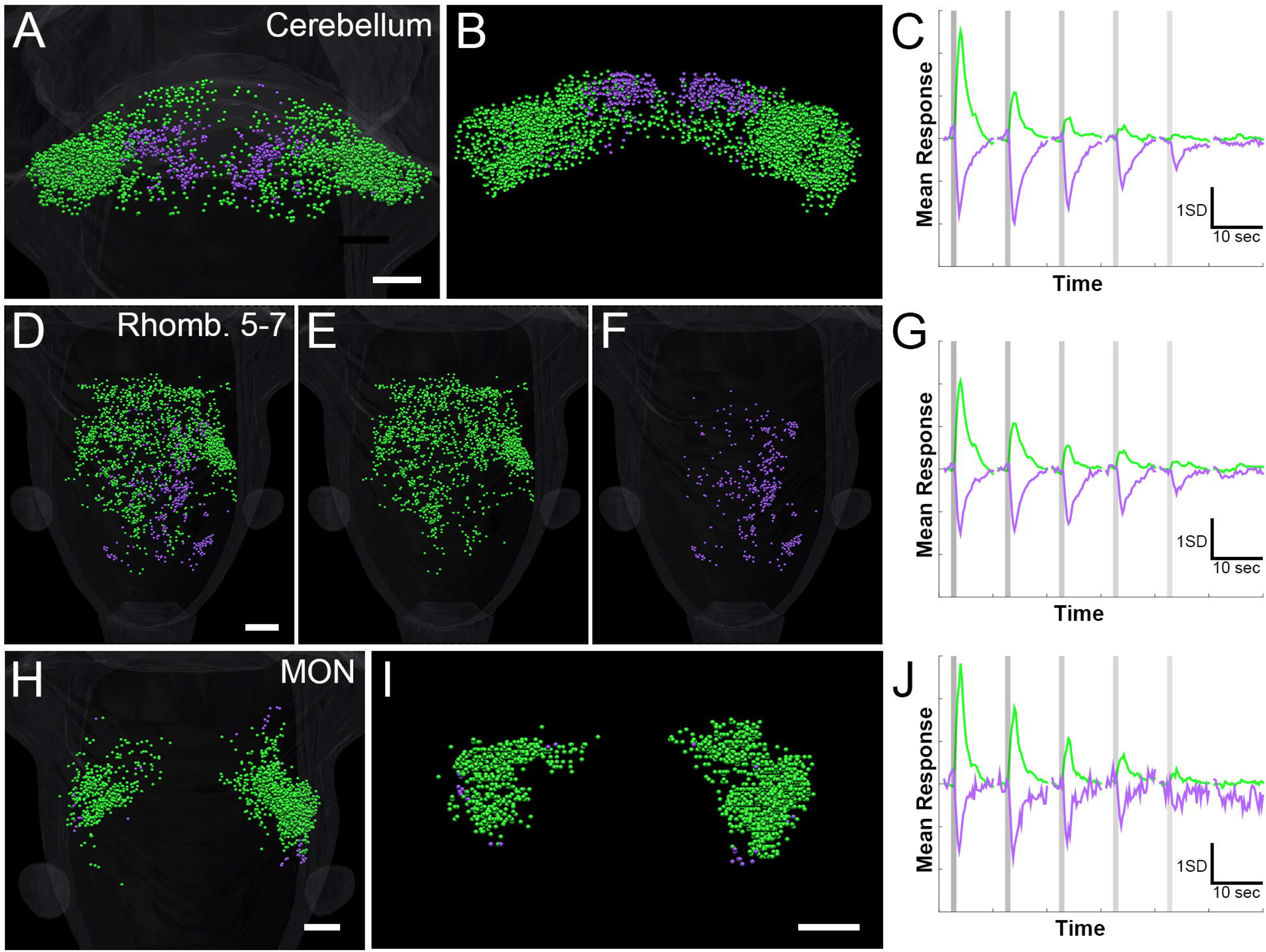
Hindbrain responses in the cerebellum, MON, and rhombomeres 5-7. Dorsal (A) and coronal (B) views of the cerebellum, showing two clusters (C) with sharp topographical distributions. Excited (D and E) and inhibited (D and F) ROIs in rhombomeres 5-7. Excited ROIs are located roughly evenly across the midline, while inhibited ROIs are dominantly ipsilateral to the stimulated (right) otolith. Average responses of the clusters present in this region are shown in (G). Dorsal (H) and coronal (I) views of the MON (clusters shown in J), where excited ROIs are bilateral, but more numerous ipsilateral to the stimulus, and a small number of inhibited neurons are scattered around the periphery. Scale bars indicate 50μm.

### Integration of bilateral vestibular information occurs at multiple locations across the brain

Applying an external force to one otolith is sufficient to elicit a tail bend in the opposite direction, and to produce a rolling motion of both eyes (ref [11] and Figure 1), so we used this simple stimulus for the basic mapping of vestibular responses described above. Our system, because we have two steerable traps, allows the independent control over the vestibular inputs to the two ears, which is difficult to achieve with natural stimuli. Using such dual OT, we next studied the neural responses to trapping of the left and right otoliths separately, and when these stimuli were combined to produce a competitive bilateral stimulus (Figure 6). With this latter stimulus, we aimed to identify regions and cellular responses involved in combining or cancelling information across the two ears.

**Figure 6:**
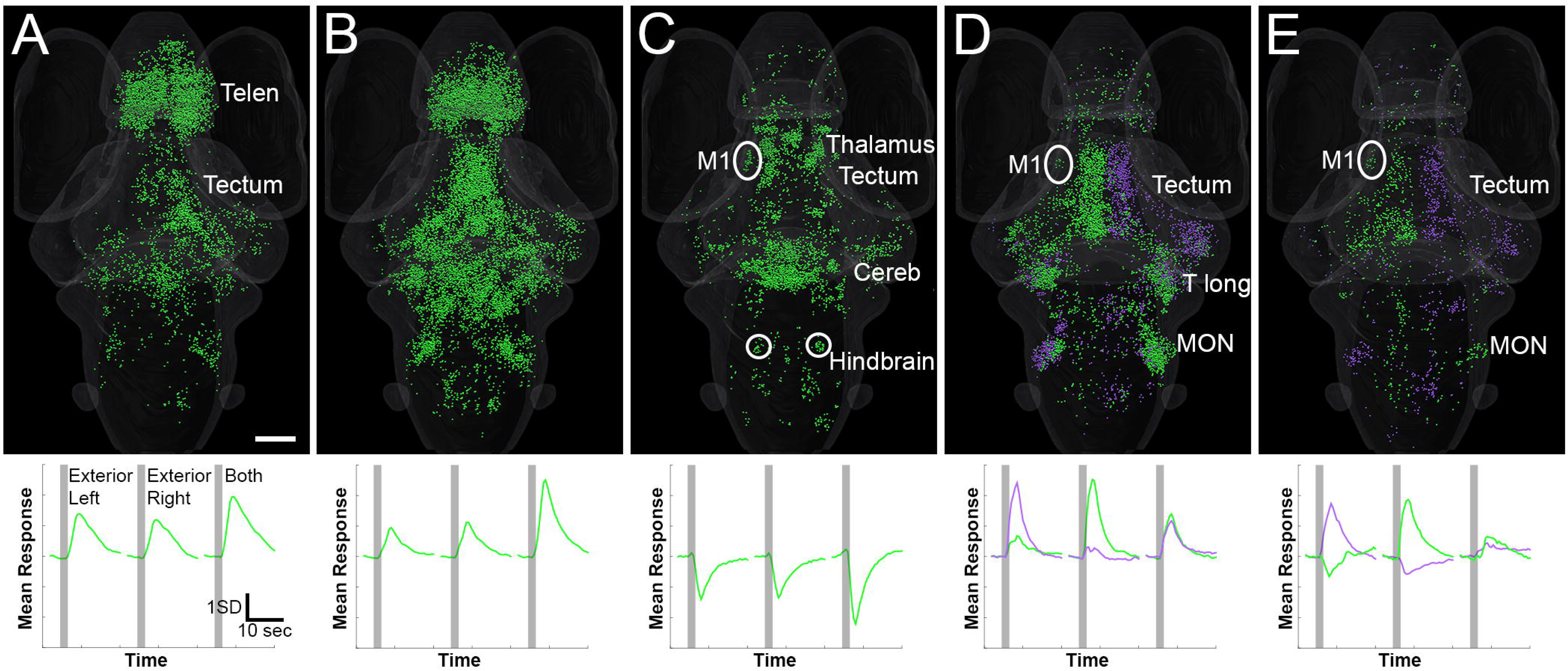
Responses to bilateral fictive vestibular stimulation. Seven functional clusters of ROIs are shown (top), each with a characteristic response profile to an optical trap on the exterior of the left otolith, the right otolith, or both traps simultaneously (bottom). Regions with notable concentrations of functional clusters are indicated, as described in the text. 3D rotations of panels A-E are shown in Supplemental Movies 4-8, respectively. Scale bar indicates 100μm, and applies to all panels. Telen: telencephalon, M1: migrated pretectal region M1, Cereb: cerebellum, T long: torus longitudinalis, MON: medial octavolateral nucleus.

One frequently occurring cluster of ROIs (Figure 6A) was excited moderately by each stimulus, with a roughly additive response to both. These ROIs were primarily located in the telencephalon, with a secondary concentration in the tectum. A second cluster (Figure 6B), scattered broadly across many regions of the brain, was principally responsive to the dual trap, with a stronger response to the combined stimulus than to the sum of the responses to the individual stimuli. Inhibited ROIs (Figure 6C) were located primarily in the cerebellum, the thalamus, the tectum, and paired nuclei in the ventral hindbrain (circles, Figure 6C), and the migrated pretectal region M1 (oval, Figure 6C-E). These ROIs were inhibited by each individual trap, with stronger responses to the dual trap. A further pair of clusters (Figure 6D) selectively responded to one of the individual traps, with a mirror-image distribution of responses to stimuli applied to the two ears. Notably, both clusters responded when both otoliths were trapped, but the magnitude of these responses was reduced as compared to those when the relevant otolith was trapped individually. The distribution of these clusters (Figure 6D) was sparse in the forebrain (although visible in the migrated pretectal region M1), generally contralateral to the stimulated ear in the midbrain (notably in the tectum and the torus longitudinalis), and principally ipsilateral to the stimulated ear in the hindbrain (especially in the MON). Bilateral cancellation was more pronounced in a further pair of clusters (Figure 6E), which were excited by stimulation of one otolith, weakly inhibited by stimulation of the other, and roughly unresponsive to the dual trap. These ROIs were notably concentrated in the tectum, although they were scattered sparsely across several other brain regions, including pretectal M1. The lateral distribution of these clusters was similar to those shown in Figure 6D, but more strictly contralateral to the stimulated ear (Figure 6E). The lone exception to this was in the MON, where responses were ipsilateral.

## Discussion

In this study, we have combined OT manipulation of the utricular otoliths with whole-brain calcium imaging to produce a brain-wide map of vestibular responses in larval zebrafish. The principal motivation for optical trapping in this setting was the ability to stimulate the vestibular system without moving the animal. It also provided the unique ability to stimulate each of the otoliths individually or in combination. Unilateral control, which can only be achieved through the removal of the otoliths in traditional vestibular experiments[33], allowed us to separate the individual contributions of the left and right otoliths to the overall neural response, while dual trapping allowed us to present conflicting bilateral stimulation that would be difficult to produce naturally. Because the semicircular canals are not yet functional in 6dpf larvae[34], and the utricular otoliths are solely responsible for detecting vestibular stimuli[35, 36], this means that we have likely stimulated the entirety of the brain’s vestibular circuitry with our traps. This has the advantage of revealing a complete and simplified version of the mammalian (or adult fish) vestibular system, reliant only on otoliths, but prevents us from addressing the mechanisms by which otolith and canal information is integrated.

The calcium imaging approach taken in this study follows in the footsteps of prior whole-brain imaging studies in zebrafish, and includes the segmentation of responses into ROIs that generally correspond to individual neurons. The goal of this was to identify not only brain regions or localized areas with responses to the stimuli, but to characterize the response profiles of the individual neurons composing these regions. Our identification of distinct functional categories of excited and inhibited neurons, often in close physical proximity, highlights how this approach may detect patterns of activity that would be overlooked or misinterpreted without segmentation.

The larval zebrafish vestibular system has previously been explored in a targeted manner, with anatomical and functional descriptions of specific sensory, sensorimotor, or motor circuits involved in detecting or responding to vestibular stimuli. The essential circuit for the VOR comprises 1) primary afferent neurons relaying information from utricular hair cells to the octavolateral neuropil in the hindbrain[37], 2) tangential neurons relaying this information to the contralateral oculomotor nucleus, and 3) the oculomotor neurons themselves[33]. Among the tangential neurons, Bianco et al [33] have identified three categories, two of which (ascending and ascending/descending) innervate the nMLF, and one of which (ascending/descending) also projects to hindbrain rhombomeres 4-8, where it is presumed to innervate reticular neurons controlling axial or fin musculature. A third category, the commissural tangential neurons, innervate the contralateral octavolateral neuropil, perhaps permitting bilateral integration of vestibular signals from the two ears[33]. At the motor level, Thiele et al [38] have identified the nMLF as sufficient for driving tail deflections like the ones that we see in response to fictive vestibular stimuli (Fig 1 and ref [11]), and Schoppik et al[39] have identified vestibular neurons in rhombomeres 5-7 that control gaze stabilization.

These detailed observations mesh with the whole-brain imaging that we have reported here. The abundant excitation that we report in the MON likely includes the tangential neurons relaying vestibular information to the oculomotor neurons to control rotation of the eyes. Our contralateral activation (and to a lesser degree, ipsilateral inhibition) of the nMLF is, presumably, a product of tangential neuron input and the cause of the tail bends that we observe during fictive stimulation. Likewise, the asymmetrical inhibition that we see in rhombomeres 5-7 may spring indirectly from descending tangential input, and may contribute to gaze stabilization. The strongly lateralized activity in the migrated pretectal region M1 likely relates to control of eye position, as neurons in this area are visually direction sensitive and involved in generating eye movements during the optokinetic response[40].

These anticipated patterns, however, represent a small portion of the responses that we observed across the brain. Analyzing neurons across all regions, we found three broad types of responses, including excitation and inhibition that were graded to the strength of the stimulus. A third category contained neurons with responses that were only weakly tuned to stimulus strength, and that maintained clear responses to the weakest stimuli. Regional analyses showed that these neurons were present only in the telencephalon and the habenulae. This localization, combined with the lack of rate coding in their responses, suggests that these neurons may encode more abstract elements of alertness or context, rather than being directly involved in the production of physical adjustments to vestibular inputs.

The rest of the brain contained ROIs with rate coded excitation and inhibition, and these responses were often lateralized. In some cases, this lateralization would have been expected, such as greater excitation in the MON ipsilateral to the stimulus, given that these neurons are the first to receive input from the sensory ganglia. Excitation in the contralateral MON, albeit at lower levels, is likely a product of communication between the two MON, possibly through commissural tangential neurons[33]. As suggested by Bianco et al[33], such contralateral communication between the MON may provide increased sensitivity to acceleration and tilt, as has also been suggested in mammals[41, 42]. As described above, the opposite lateralization of nMLF responses is predicted based on the nMLF neurons’ established role in producing postural adjustments in the tail[38], which bends contralaterally from the stimulus[11]. Across other brain regions, there is a trend toward greater excitation on the contralateral side (as seen in the thalamus, tegmentum, and rhombomeres 5-7), and greater inhibition on the ipsilateral side (seen clearly in the thalamus, strikingly in the tectal PVL and neuropil, and also in rhombomeres 5-7). These results for the tectum, thalamus, and tegmentum are difficult to put into a functional context, given the lack of knowledge about the functional connectivity of these brain regions in larval fish. Asymmetrical inhibition in rhombomeres 5-7 may subserve forward swim bouts that correct for pitch instability[43] and the asymmetrical recruitment of spinal circuits involved in roll correction[44]. Finally, responses in the cerebellum are bilaterally symmetrical, and are notable for the tight restriction of inhibition to a small region, likely of Purkinje cells, near the dorsal midline. This region has been shown to send bilateral projections to other regions that we have shown to be responsive to vestibular stimuli, including the nMLF, thalamus, and reticular formation[45]. This places localised inhibition of Purkinje cells, and associated disinhibition of the output eurydendroid cells, in a position where it could modulate behavioral responses to vestibular stimuli.

In dual trap experiments, we explored interactions of information from the two ears. (Figure 6). Many of the responses to simultaneous traps of both otoliths were enhanced versions of the responses to traps of one otolith or the other (Figure 6A-C), that is to say that they were combining the inputs in an additive or superadditive manner. ROIs summing the inputs in a roughly additive manner (Figure 6A) were prominently (but not exclusively) located in the telencephalon. These may be the same ROIs that, in variable-power trap experiments of individual otoliths, showed relatively power-insensitive responses. Those that were excited in a superadditive manner (Figure 6B) showed a broader distribution, reflecting excitation observed throughout many brain regions in the prior experiments (Figures 2-5). Inhibited ROIs (Figure 6C) were distributed similarly to those previously observed (Figures 2-5), with notable concentrations in the cerebellum, thalamus, tectum, and small paired nuclei in the ventral hindbrain. These three clusters of ROI have in common that they are equally sensitive to external forces on both otoliths, suggesting that they report on the presence of, but not the mechanical details of (and therefore the specific simulated vestibular inputs corresponding to) the different fictive stimuli. This is in contrast to the final two pairs of clusters (Figure 6D and E), where ROIs show preferential responses to one unilateral stimulus (Figure 6D) or opposing responses to the two unilateral stimuli (Figure 6E). In these cases, the dual trap attenuated or eliminated the responses that were observed to the preferred individual stimulus. This suggests that these ROIs are receiving input from both ears, and that the non-preferred ear’s signals are capable of offsetting the preferred ear’s normal responses when they contain contradictory vestibular information. These clusters’ most prominent concentrations are the diencephalon and midbrain, with few ROIs in the telencephalon, and relatively few in the hindbrain (excepting the MON, Figure 6D and E). This suggests that the bilateral interactions may take place at the stage of vestibular processing and perception, rather than through gating in hindbrain motor circuits, although our results do not exclude the possibility of small populations of hindbrain neurons that directly modulate behaviors based on conflicting signals from the two ears.

Across all of these observations, the breadth and richness of the responses suggests a more extensive system for vestibular processing across larval zebrafish brain than has previously been appreciated. These results highlight both the strengths and the limitations of our whole-brain imaging approach. The results of past studies on specific circuits provide ground truths to suggest that our imaging is accurately capturing vestibular processing. The results of our comprehensive and unbiased observations reveal a host of additional regional responses, distinct response types, topography, and bilateral asymmetries that may have important implications for how the brain processes vestibular information. These results do little, however, to elucidate the circuit-level mechanisms mediating these responses. Rather, they provide a departure point for more targeted studies into these neurons’ anatomy, connectivity, neurotransmitter use, and behavioral contributions.

## Acknowledgements

We thank members of the Scott and Rubinsztein-Dunlop labs for feedback on the manuscript. Support was provided by the an NHMRC Project Grant (APP1066887), ARC Future Fellowship (FT110100887), a Simons Foundation Explorer Award (336331), and two ARC Discovery Project Grants (DP140102036 & DP110103612) to E.K.S.; an ARC Discovery Project (DP140100753) to H.R-D.; and a fellowship from the Human Frontiers Science Program (LT000146/2016) to M.A.T.

## Author Contributions

I.A.F, H.R.-D. and E.K.S conceived the project. I.A.F performed experiments. G.V. performed bioinformatics analyses. I.A.F. and M.A.T. designed and built the apparatus. I.A.F, G.V, M.A.T, H.R. and E.K.S and wrote the manuscript.

## STAR METHODS

### CONTACT FOR REAGENT AND RESOURCE SHARING

Further information and requests for resources, reagents, or data should be directed to and will be fulfilled by the Lead Contact, Ethan Scott.

### EXPERIMENTAL MODEL AND SUBJECT DETAILS

#### Animals

All procedures were performed with approval from The University of Queensland Animal Welfare Unit (in accordance with approval SBMS/378/16). Zebrafish (*Danio rerio*) larvae, of either sex, were maintained at 28.5°C on a 14hr ON/10hr OFF light cycle. Adult fish were maintained, fed, and mated as previously described [46]. All experiments were carried out in *nacre* mutant *elavl3:H2B*-*GCaMP6s* larvae of the TL strain.

### METHOD DETAILS

#### Sample preparation

Zebrafish 6 day post-fertilization (dpf) larvae were immobilised dorsal side up in 2% low melting point agarose (Sigma-Aldrich) on microscope slides. The agarose on the rostral and right side of the fish was cut vertically, parallel to the fish, with a scalpel, and removed. The embedded fish was transferred to custom made, 3D printed chamber. The agarose surrounding the tail was freed by removing segments of agarose perpendicular to the tail until reaching the swim bladder. The chamber was filled with E3 media. Larvae were then transferred to the imaging room and allowed to acclimate for 15 min prior to imaging on the custom-built dual optical trapping microscope presented in Figure 1.

#### Dual Optical Trapping (OT) system

The dual OT system (Figure 1) was composed of a IR laser (1070nm IPG Photonics YLD-5 fibre laser), a half wave plate that rotates polarisation by 45 degrees, and a polarising beam splitter that splits the incoming beam into two beams of same intensity with perpendicular polarisation. The two independent beams were reflected off of gimbal mirrors (GM) (Thorlabs GM100). The two beams were recombined with a second polarising beam splitter and a telescope with lenses L1 (150 mm focal length,) and L2 (300 mm focal length). The beams were then reflected off of a 950 nm cut-off wavelength shortpass dichroic mirror in the imaging column, and projected onto the back focal plane of a 20x 1NA Olympus microscope objective (XLUMPLFLN-W). This created two tightly focussed spots at the imaging plane of the microscope objective. The positions (x,y) of each spot were steered with the GMs. Two shutters (S1 and S2, Thorlabs SHB1T) allowed independent gating of the OT and were driven using an Arduino (Leonardo). Laser power intensities (0, 100, 200, 300, 450 and 600 mW) were gauged using a power meter at the focal plane of the 20x 1NA objective.

#### Fluorescence imaging system

Calcium imaging and measurements of eye movement were performed through the same 20x objective that was delivering the OT. For the imaging of a chosen depth on the PCO edge 5.5 camera (Camera 1), a combination of filter (F, Thorlabs FF01-517/520-25), tube lens (L3, 180mm focal length, Thorlabs AC508-180-A), relay lenses (Lr, Thorlabs AC254-125-AML), ETL (Optotune EL-10-30-Ci-VIS-LD driven with Gardasoft TR-CL180) and offset lens (Lo, Eksma Optics 112-0127E) was constructed as described by Fahrbach et al[47]. With this configuration, we were able to scan 300μm of brain tissue above the original imaging plane, where the OT is applied.

The scanning light sheet was generated using a 488nm laser (OBIS 488 lx) combined with 2D galvo mirrors (Thorlabs GVSM002/M), a 50/50 beamsplitter, and a 1D line diffuser (RPC Photonics EDL-20-07337). One galvo scanning direction (Y) created the light sheet while the second direction (Z) created the depth scan in the sample. The two mirrors were driven independently using Arduinos (DUE) with custom-written code. The Y scanning was a sawtooth scan at 600 Hz, which was synchronized to the camera acquisition to ensure similar illumination for each camera acquisition. The Z galvo was driven in steps to scan the light-sheet through the sample. The 50/50 beamsplitter created two light sheets, one projecting into the rostral side and one into the right side of the fish. A 1D line diffuser was placed just after the galvos to reduce shadowing in the planes.

In our experiments, an exposure time of 10ms was chosen for each plane during volumetric imaging, with laser power output of 60mW. A total Z scan of 250 μm was performed with 10 μm steps which resulted in a 4Hz brain scanning speed.

#### Behavioural imaging system

To image the whole larva and record its tail movements, a 4x 0.1NA Olympus microscope objective (PLN 4X) was placed below the sample chamber, and a tube lens L4 projected onto a Basler aca1920 camera (Camera 2) recording movements at 30 fps. Note that the 4x and 20x objectives are not collinear.

### QUANTIFICATION AND STATISTICAL ANALYSIS

#### Measuring tail and eye movements

Since whole-brain imaging was performed at 4Hz, the eye movements were also measured at this speed. Tail movements were recorded at 30Hz using Camera 1.

Tail bends were imaged along the length of the tail, and clearly visible locations on the tail were tracked horizontally (perpendicular to the tail) using Gaussian fitting in Matlab. The angle of deflection was calculated with simple trigonometry by setting the base of the tail as a vertex, the initial position of the tracked marker as one point, and the moving marker as the other point.

Eye motions were tracked in the same dorsal-ventral plane as the utricular otoliths. Visible landmarks on the pigmented retinal epithelium were used to track consistent locations. The displacements of these landmarks were measured by recording center points of the landmarks, and the value for a given eye in a given trial was the average movement of the landmarks on that eye. The same Matlab code used for the tail was used for the eyes. Roll angles were calculated with simple trigonometry, using 300μm as the average height of 6dpf larvae’s eyes, and assuming rotation around the centre of the eye.

#### Extraction of fluorescent traces

We used the CaImAn package to analyse our images [23] (github.com/flatironinstitute/CaImAn). The movies were first motion corrected [48], then the fluorescent traces were demixed and denoised with 4000 components per slice [23].

#### Whole brain analysis of fluorescent traces

The resulting ROIs and fluorescent traces were further analysed in Matlab. The traces from the thirteen fish were pooled and z-scored. For the first set of experiments, predictors were built for each of the stimulus onset and offset, with a typical GCaMP response occurring for each of the three presentations of the tone. The coefficient of determination (r^2^) of the linear regression models was used to select stimulus responsive ROIs, and we chose a 0.1 threshold based on the r^2^ distribution of our models to allow for conservative filtering of the data.

After this filtering step, the traces from ROIs deemed responsive to the stimuli were clustered using k-means with the cityblock distance and three replicates. The clusters were then compared to the predictors using linear regression and the clusters showing r^2^ values above 0.4 were kept for further analysis.

#### Registration of the results to a reference brain

We used Advanced Normalization Tools (ANTs, github.com/ANTsX/ANTs) to register our results on the H2B-RFP reference of Zbrain [24, 26, 49]. The time averaged movie stacks were used to build a common template, before registering this template to the Zbrain atlas (See Table S1 for the ANTs calls). The resulting warps were sequentially applied to the centroids of extracted ROIs to map them all in the same frame of reference.

The Warped ROI coordinates were then placed in each of the 294 brain regions defined in the Zbrain atlas [26].

#### Analysis of individual brain regions

The ROI traces belonging to ROIs from the 11 analyzed brain regions, as defined in the Zbrain atlas, were analyzed as the whole brain data.

#### Data visualization

We used Unity^®^ to represent each ROI centroid as a sphere of diameter 4. An isosurface mesh of the zebrafish brain was generated from the Zbrain masks for Diencephalon, Mesencephalon, Rhombencephalon, Telencephalon and the eyes using ImageVis3D [50]. The mesh was imported in Unity^®^ and overlaid to the ROIs.

### DATA AND SOFTWARE AVAILABILITY

All data and software will be made available upon request.

#### Primary Supplemental PDF

**TABLE S1, Key Resource Table:** The identities of the calls used in ANTs registration.

Primary Supplemental PDF:
|  |  |
| --- | --- |
| Template building | antsMultivariateTemplateConstruction2.sh -d 3 -b 1 -c 2 -g 0.1 -j 16 -i 10 -r 1 -n 0 -m CC -t SyN -o \$Template *.tif |
| Registration | antsRegistrationSyNQuick.sh -d 3 -o \$TemplateToZbrain -f \$ZBrain_template -m \$Template |
| Centroids warp | antsApplyTransformsToPoints -d 3 -i \$ROIs -o \$Warped_ROIs -t [\$TemplateToZbrainAffine, 1] -t \$TemplateToZbrainInverseWarp -t [\$FishToTemplateAffine.mat, 1] -t \$FishToTemplateInverseWarp |

**Supplemental Movie 1: Tail movements elicited by OT**

Videos are shown as optical traps of different powers are applied to the lateral edge of the left otolith. The head of this animal is embedded in agarose, but the tail is free, and the tail’s movements to the optical trap are clearly observable. The flickering in the head is a result of the volumetric SPIM.

**Supplemental Movie 2: Eye movements elicited by OT**

Videos are shown as optical traps of different powers are applied to the lateral edge of the left otolith. The trapping laser is indicated by the red circle in the bottom left of the videos. Subtle rotational eye movements can be observed, especially with stronger traps.

**Supplemental Movie 3: 3D rotation of brain-wide responses to OT**

This is a 3D movie of ROIs belonging to three functional clusters, registered against the Z-brain atlas. Clusters are color-coded, as described in Figure 2.

**Supplemental Movie 4: 3D rotation of Figure 6A**

A 3D rotation of ROIs excited by external traps of either otolith, for which the response to a dual trap is roughly the linear summation of the individual responses.

**Supplemental Movie 5: 3D rotation of Figure 6B**

A 3D rotation of ROIs excited by external traps of either otolith, for which the response to a dual trap is greater than that of the combined individual responses.

**Supplemental Movie 6: 3D rotation of Figure 6C**

A 3D rotation of ROIs inhibited by external traps of either otolith.

**Supplemental Movie 7: 3D rotation of Figure 6D**

A 3D rotation of ROIs excited by external traps of either the left or the right otolith. The responses to dual traps are an attenuated version of the response to the preferred otolith alone, as described in Figure 6D. Green ROIs are responsive to traps of the right otolith, and purple ROIs are responsive to traps of the left otolith.

**Supplemental Movie 8: 3D rotation of Figure 6E**

A 3D rotation of ROIs excited by external traps of either the left or the right otolith, and inhibited by traps of the non-preferred otolith. The responses to dual traps are minimal, as shown in Figure 6E. Green ROIs are excited to traps of the right otolith, and purple ROIs are excited to traps of the left otolith.

